# Mnemonic prediction errors bias hippocampal states

**DOI:** 10.1101/740563

**Authors:** Oded Bein, Katherine Duncan, Lila Davachi

## Abstract

In situations when our experience violates our predictions, it is adaptive to upregulate encoding of novel information, while down-weighting retrieval of erroneous memory predictions to promote an updated representation of the world. We asked whether mnemonic prediction errors promote distinct hippocampal processing ‘states’ by leveraging recent results showing that encoding and retrieval processes are supported by distinct patterns of connectivity, or ‘states’, across hippocampal subfields. During fMRI scanning, participants were cued to retrieve well-learned room-images and were then presented with either an image identical to the learned room or a modified version (1-4 changes). We found that CA1-entorhinal connectivity increased, and CA1-CA3 connectivity decreased, with the number of changes to the learned rooms. Further, stronger memory predictions measured in CA1 during the cue correlated with the CA1-entorhinal connectivity increase in response to violations. Our findings provide a mechanism by which mnemonic prediction errors may drive memory updating - by biasing hippocampal states.

## Introduction

As our day unfolds, much of what we encounter is expected: we typically navigate to work or school along the same route, sit in the same seats in the same space and engage with the same people. However, layered on top of the repetition of similar places and events are novel or surprising events; and when we travel to unfamiliar places, we experience even more novelty. This interplay between similarity and novelty poses different demands on our memory system. On the one hand, the repeating aspects of each day may trigger the retrieval of related memories that may allow those memories to then serve as predictions to guide adaptive behavior (Bar, 2009; Lisman & Redish, 2009; Stachenfeld, Botvinick, & Gershman, 2017). By contrast, surprising events may shift the memory system towards encoding of those contextually novel events (Duncan, Sadanand, & Davachi, 2012; Hasselmo & Stern, 2014; Hasselmo, Wyble, & Wallenstein, 1996; Kumaran & Maguire, 2007b; Meeter, Murre, & Talamini, 2004). Intriguingly, the hippocampus has been proposed to mediate both the encoding of new events and the retrieval of previous related experiences (Eichenbaum, Yonelinas, & Ranganath, 2007; Marr, 1971; Scoville & Milner, 1957; Squire & Alvarez, 1995). However, at a mechanistic level, these processes require seemingly conflicting processes: new encoding benefits from plasticity in hippocampal networks while this kind of plasticity during retrieval may permanently alter the veracity of long-term memories (Hasselmo, Bodelón, & Wyble, 2002; Hasselmo & Stern, 2014; O’Reilly & McClelland, 1994). Furthermore, at the neural population level, encoding presumably requires that current experiences be represented in an activity pattern distinct from other stored memories, a process known as ‘pattern separation’ (O’Reilly & McClelland, 1994; Yassa & Stark, 2011). Retrieval, on the other hand, may be supported by the recovery of a previously encoded activity pattern, or ‘pattern completion’ (Knierim & Neunuebel, 2016; Marr, 1971; O’Reilly & McClelland, 1994; Treves & Rolls, 1994). Thus, a critical question is how can the hippocampal system balance these two seemingly opposing processes? And what factors may bias the hippocampus towards one over the other? (Colgin, 2016; Colgin et al., 2009; Duncan, Sadanand, et al., 2012; Duncan, Tompary, & Davachi, 2014; Hasselmo & Stern, 2014; Hasselmo et al., 1996).

Current models of hippocampal function propose that communication along distinct CA1 pathways may be associated with encoding and retrieval ‘states’ (Colgin, 2016; Hasselmo et al., 2002; Hasselmo & Stern, 2014). Specifically, it has been proposed that, during encoding of novel experiences, input from the medial temporal cortical regions that receive numerous sensory inputs such as the entorhinal cortex (Burwell, 2000; McClelland, McNaughton, & Oreilly, 1995; Schultz, Sommer, & Peters, 2015; Suzuki & Amaral, 1994), may be prioritized by hippocampal area CA1. By contrast, during retrieval, CA1 may preferentially process input from hippocampal area CA3. CA3 neurons are highly-interconnected, a feature proposed to facilitate pattern completion and promote the retrieval of encoding related ensembles, which can then be conveyed to area CA1 (Marr, 1971; Montgomery & Buzsaki, 2007; Nakazawa et al., 2002; Norman & O’Reilly, 2003; O’Reilly & McClelland, 1994; Rolls, 2016; Treves & Rolls, 1994). Empirical work in rodents has shown that CA1-entorhinal coherence is higher in the fast-gamma band compared to the slow-gamma band, while CA1-CA3 coherence is higher in the slow versus fast gamma band, supporting a functional distinction between these pathways (Colgin et al., 2009; Kemere, Carr, Karlsson, & Frank, 2013). These different gamma band frequencies have also been linked to different behaviors such as fast or slow running speed (Colgin, 2016; Kemere et al., 2013; Zheng, Bieri, Trettel, & Colgin, 2015). More recently, CA1 fast-gamma band activity was observed during learning of spatial routes in a maze, compared to slower-gamma activity evident during retrieval of learned routes (Lopes-dos-Santos et al., 2018). Further, CA1-CA3 coherence has been shown to be enhanced in the central arm of a T-maze, potentially reflecting retrieval of the goal location (Montgomery & Buzsaki, 2007). Additional support of the dissociation between the two pathways comes from studies showing that CA1 coupling with entorhinal cortex and area CA3 occurs at different phases of a theta cycle (Fernández-Ruiz et al., 2017; Hasselmo et al., 2002; Hasselmo & Stern, 2014; Newman, Gillet, Climer, & Hasselmo, 2013; Schomburg et al., 2014; Tort, Komorowski, Manns, Kopell, & Eichenbaum, 2009). Extending this theoretical and empirical framework to humans, we have recently shown, using functional magnetic resonance imaging (fMRI), that CA1-CA3 functional connectivity is significantly enhanced during episodic memory retrieval compared to novel associative encoding (Duncan et al., 2014). Importantly, the magnitude of CA1-CA3 connectivity during retrieval predicted retrieval success (Duncan et al., 2014). Together, these results provide support for the idea that the hippocampus may shift between encoding and retrieval ‘states’ by modulating CA1 connectivity with distinct input regions.

One prominent factor that may bias hippocampal dynamics towards encoding rather than retrieval is mnemonic prediction error (Hasselmo, Schnell, & Barkai, 1995; Hasselmo et al., 1996; Meeter et al., 2004). There is now much work demonstrating that hippocampal activity increases when sequential predictions are violated (Axmacher et al., 2010; Chen, Cook, & Wagner, 2015; Kumaran & Maguire, 2006, 2007a). This increase has been localized to hippocampal area CA1 in both humans and rodents (Allen, Salz, McKenzie, & Fortin, 2016; Chen et al., 2015; Chen, Olsen, Preston, Glover, & Wagner, 2011; Duncan, Ketz, Inati, & Davachi, 2012). One interpretation of this increased CA1 BOLD signal during mnemonic prediction errors is that it may facilitate the encoding of the novel, unexpected, information, and thus promote memory updating and the improvement of future predictions (Henson & Gagnepain, 2010; McClelland et al., 1995). Indeed, there is some behavioral evidence that mnemonic prediction errors facilitate episodic memory (Chen et al., 2015; Greve, Cooper, Kaula, Anderson, & Henson, 2017). We set out examine whether mnemonic prediction errors are associated with a shift in hippocampal processing towards an encoding state that prioritizes input from entorhinal cortex and away from a retrieval state (Colgin & Moser, 2010; Hasselmo & Stern, 2014; Meeter et al., 2004; O’Reilly & McClelland, 1994). Furthermore, we aimed to link these effects with the quality of the prediction itself.

To test these hypotheses, participants underwent extensive training to learn the furniture and layout of 30 distinct rooms. Then, in the fMRI scanner, we probed participants to retrieve each learned room by presenting a verbal cue (e.g. Johnsons boy’s bedroom), which was then followed by a room image that either matched the learned room image or included changes (Figure 1A). We operationalized the retrieval of the image as a form of memory ‘prediction’ and prediction errors were cases when the presented perceptual image was a violation of the actual learned image. Using high resolution imaging, we find that mnemonic prediction errors biased CA1 functional connectivity towards entorhinal cortex and away from subregion CA3. Moreover, the extent to which the hippocampus exhibited a shift into an encoding state during mnemonic prediction errors correlated with the strength of the prediction. Taken together, these findings show that mnemonic prediction errors bias CA1 functional connectivity, potentially to shift hippocampal processing to favor encoding and down-weight retrieval.

**Figure 1.**
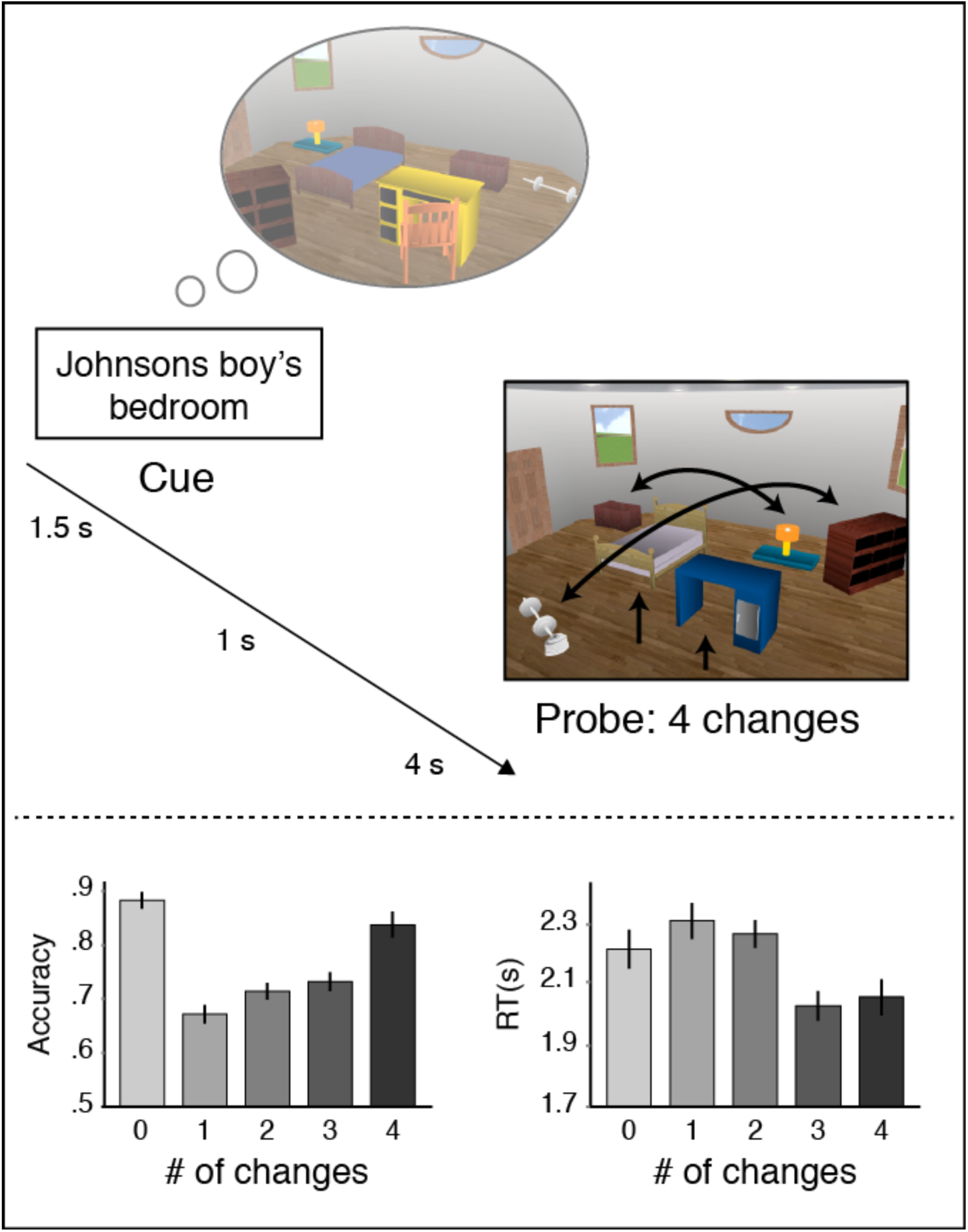
Top: Trial example: participants were presented with a cue probing them to retrieve a room image that they had extensively learned prior to the scan. After a short delay, they saw a probe image that included 0-4 changes relative to the learned image (4 changes here), and indicated whether the seen image matched the learned image (see Methods). Bottom right: accuracy and reaction times (RTs) in the match task.

## Results

### Behavior

A full reporting of the behavioral results has been provided in Duncan et al. (2012) and is summarized here following a brief description of the task. We had 2 types of change-detection task: a Furniture task and a Layout task, in which participants indicated whether a change occurred in the identity or the layout of the furniture, correspondingly. On each trial, the room image included 0-2 task relevant changes and 0-2 irrelevant changes. For example, in the Furniture task there can be 2 task-relevant changes in the identity of the furniture, and 1 task-irrelevant change in the layout of the furniture (see Methods). As reported in Duncan et al. (2012), A 2 (Task) by 3 (Relevant changes) by 3 (Irrelevant changes) repeated-measures ANOVA revealed that participants were more accurate in the Layout task compared to the Furniture task. Relevant changes did not interact with Task, however, introducing irrelevant changes did reduce accuracy in the Furniture task more than in the Layout task. Finally, relevant and irrelevant changes interacted, such that having no irrelevant changes increased accuracy, but only if there were no relevant changes as well (for more details, see Duncan et al., 2012). However, despite some differences in behavioral effects of irrelevant and relevant changes, CA1 BOLD response predominately tracked the total number of changes, irrespective of relevance to the task (Duncan et al., 2012). Thus, in subsequent analyses we collapse across relevant and irrelevant changes and report the behavioral and neural data as a function of the total number of changes. Accuracy data in the change-detection tasks were entered to a 5 (Changes: 0-4) by 2 (Task: Furniture/Layout) repeated measures ANOVA. This ANOVA revealed main effects of Changes and Task, as well as an interaction (Changes: *F*(4,72) = 33.48, *p* < .001; Task: *F*(1,18) = 8.50, *p* < .01; Interaction: *F*(4,72) = 3.24, *p* < .02). In both tasks, accuracy was highest when there was no change (0-change) and in the 4-changes conditions in comparison to the 1- to 3-changes conditions.

Response times (RTs) also tracked the accuracy data: RTs were significantly shorter in the 0-changes and the 4-changes conditions compared to the 1- to 3-changes. These results reflect the relative ease of indicating “match” when there were no changes at all, or “mismatch” when there were many changes which provides support for the rooms having been well learned. RTs were also entered into the same ANOVA as the accuracy data, which again revealed main effects of Changes and Task, and an interaction (Changes: *F*(4,72) = 5.83, *p* < .001; Task: *F*(1,18) = 7.57, *p* < .02; Interaction: *F*(4,72) = 9.1, *p* < .001). Mean and SD of accuracy and RT in each of the number of changes and each task are provided in Table 1, and collapsed across tasks in Figure 1. Importantly, in the neural data we did not observe a main effect of Task nor an interaction between Task and Changes; thus, we collapsed across tasks (see Results).

**Table 1.** Accuracy rates and reaction times in the Layout and Furniture tasks. Reaction times are in seconds. SDs are in parentheses

| <u>Accuracy:</u> |  |  |  |  |  |
| --- | --- | --- | --- | --- | --- |
| Task | Number of changes |  |  |  |  |
|  | 0 | 1 | 2 | 3 | 4 |
| Layout | .89 (.10) | .69 (.09) | .74 (.08) | .75 (.10) | .90 (.11) |
| Furniture | .88 (.10) | .65 (.10) | .68 (.10) | .72 (.11) | .77 (.17) |
| <u>Reaction times:</u> |  |  |  |  |  |
| Task | Number of changes |  |  |  |  |
|  | 0 | 1 | 2 | 3 | 4 |
| Layout | 2.29 (.34) | 2.36 (.30) | 2.37 (.27) | 2.27 (.32) | 2.04 (.37) |
| Furniture | 2.30 (.35) | 2.42 (.30) | 2.40 (.27) | 2.28 (.29) | 2.37 (.34) |

### Mnemonic prediction errors decrease CA1-CA3 functional connectivity, while increasing CA1-Entorhinal connectivity

Functional connectivity was measured using a beta-series correlation approach (Rissman, Gazzaley, & D’Esposito, 2004). Prior to testing our main hypothesis, we conducted, in each pair of anatomically defined ROIs, a 5 (Changes: 0-4) by 2 (Task: Furniture/Layout) repeated-measures ANOVA, to test whether collapsing across tasks is warranted. Indeed, there was no main effect of Task nor a Changes by Task interaction in functional connectivity between CA1-CA3 (The CA3 ROI included CA2,CA3, and dentate gyrus) or CA1-entorhinal, for the left and the right hemispheres (all *p*’s > .17). Given this, we collapsed across tasks for our main analyses. In the left hemisphere, we found an interaction between Changes (0-4) and ROI (entorhinal, CA3) using a repeated measures ANOVA (*F*(4,72) = 6.04, *p* < .001, η*p*^2^ = 0.25), confirming our prediction that the number of changes in the presented room differentially modulated CA1 connectivity with entorhinal cortex and area CA3 (Figure 2). However, the same ANOVA conducted on the right hemisphere did not reveal a significant interaction (*p* > .68; no main effect of Changes, *p* > .96; a main effect of ROI was observed, *p* < .005). This laterality of the interaction was also confirmed by a 3-way interaction of Hemisphere (right, left) by ROI (CA3, entorhinal), by Changes (0-4) (*F*(4,72) = 4.24, *p* < .005, η*p*^2^ = 0.19). Thus, due to the specificity of the interaction to the left hemisphere, further analyses were restricted to the left hemisphere ROIs.

**Figure 2.**
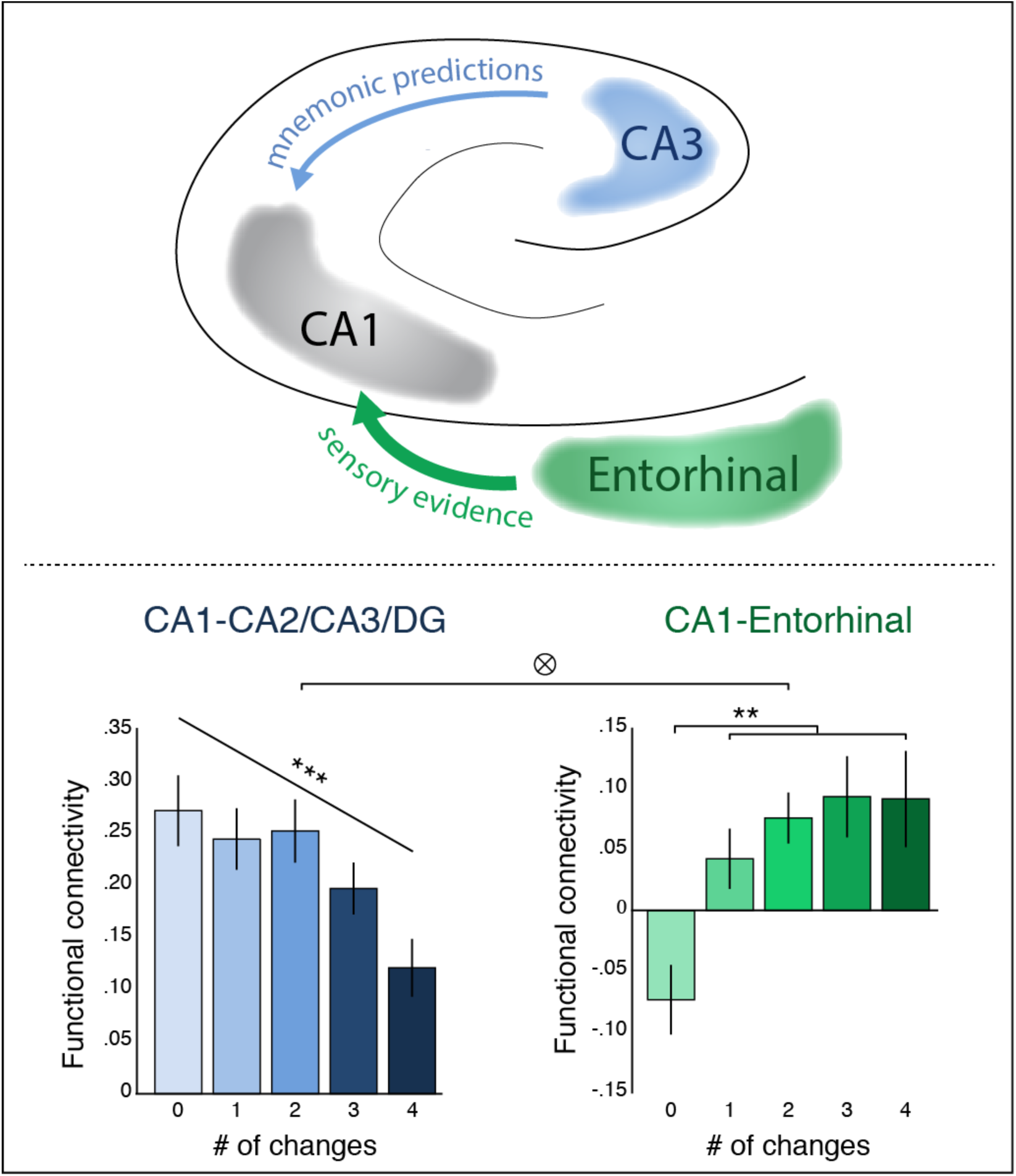
Functional connectivity with region CA1. Top: mnemonic prediction errors decreased CA1-CA3 interaction, while increasing CA1-Entorhinal cortex interaction, potentially reflecting reduced processing of erroneous predictions, and up-regulating processing of sensory evidence. Bottom: functional connectivity of CA1 with region CA2/CA3/DG (blue) and Entorhinal (green), for each number of changes. F-transformed beta-series correlation was our measure of functional connectivity. Data are from the left hemisphere (see main text). ** p < .01, *** p < .005

Having established that the number of changes differentially modulated connectivity in CA1 pathways, we moved on to examine the connectivity of CA1 with each region (entorhinal, CA3) separately. As predicted, a one-way ANOVA with the factor of Changes (0-4) revealed a significant *increase* in CA1-entorhinal connectivity as number of changes increased (*F*(4,72) = 4.49, *p* < .003, η*p*^2^ = 0.20). By contrast, and again consistent with our predictions, CA1-CA3 connectivity *decreased* as number of changes increased (*F*(4,72) = 3.58, *p* < .02, η*p*^2^ = 0.17).

Although not the main aim of the current study, we sought to further characterize the observed connectivity changes. To that end, we asked, for each pair of ROIs (CA1-entorhinal/CA1-CA3), whether connectivity changes correspond more to a linear trend, or rather to a simpler match < mismatch pattern. For each pair of ROIs, we constructed a mixed-level model in which functional connectivity was the explained variable. As explaining variables, we included both a linear trend contrast in which the number of change (0-4) were coded as linearly increasing numbers, and a match < mismatch contrast, in which the 0-change condition (i.e., match to the learned image) was compared to the 1-4 changes conditions grouped together, treating all trials with any change identically (see Methods). We then compared this full model to either a model including only the linear trend contrast, or only the match < mismatch contrast. In CA1-entorhinal connectivity, we found that the full model significantly outperformed the linear model (*χ*^2^ = 4.39, *p* < .05), but not the match < mismatch model (*χ*^2^ = 1.31, *p* > .25), suggesting that the match < mismatch contrast better describes CA1-entorhinal connectivity. For CA1-CA3 connectivity, the full mode significantly outperformed the match < mismatch model (*χ*^2^ = 8.63, *p* < .005) but not the linear model (*χ*^2^ = .59, *p* > .4), suggesting that CA1-CA3 connectivity may decrease linearly as number of changes increase.

### Functional connectivity between CA1 and entorhinal cortex correlates with mnemonic prediction strength

In the previous analysis, we operationalized mnemonic prediction error as increasing with the number of changes present in the probe room image. However, if the response is related to *prediction error*, per se, it should be modulated by the strength with which an individual uses the cue to internally generate the memory-based prediction. While participants were all extensively trained on all 30 rooms in the experiment, we could ask whether variance in mnemonic reinstatement across individuals correlates with CA1 connectivity in response to room alterations. To that end, we assessed whether the strength of the prediction, as estimated by the level of neural pattern similarity between a retrieved memory for a room compared to viewing of the same room, was related to the changes in connectivity between CA1 and entorhinal cortex or CA3 during prediction violations. Specifically, prediction-strength was estimated by correlating the multivariate BOLD activity pattern in CA1 during the presentation of each cue (e.g. ‘Johnson’s boy’s bedroom,’ to which participants were instructed to retrieve a memory of that room) with the activity pattern measured when participants actually viewed the same room (the 0-changes image of the corresponding room) and comparing it to the correlation with the pattern evoked by 0-changes images of other rooms. Thus, the strength of mnemonic prediction should be reflected by the degree to which cue periods (when memories are generated) are more correlated with viewing the same as compared to other rooms. This analysis was restricted to the left hemisphere, where we had already obtained significant connectivity differences with the number of changes (Figure 2). While we found that the correlation with the corresponding room was numerically higher than to the other rooms, the difference did not reach statistical significance (match: *M* = .004, *SD* = 0.01; other: *M* = .0004, *SD* = .005; *t*(18) = 1.3, *p* = .11, one-tailed), suggesting large variance in reinstatement. Thus, in order to ask whether individual differences in prediction strength relate to increases in CA1-entorhinal connectivity in response to the altered room images, for each subject we took the match < mismatch contrast score (0-changes vs. all levels of changes) because this score best characterized increases in CA1-entorhinal connectivity when viewing altered rooms in our experiment. Second, taking a within-participant difference score rather than a raw connectivity measure ensured that we are not simply using some baseline measure of participants’ connectivity but rather a within-participant measure of how much connectivity increased across experimental conditions. We found that prediction-strength in CA1 and the increase in CA1-entorhinal connectivity were significantly correlated (*Pearson’s r* =.51, *p* <.013, one-tailed; Figure 3), lending further support for our suggestion that functional connectivity increases are related to predictions and their violations. Prediction strength did not correlate with CA1-CA3 decreases in connectivity (linear decrease score, better accounting for connectivity changes between CA1 and CA3: *r* = .16, *p* = .74; match > mismatch score: r = −.12, p = .68; one-tailed).

**Figure 3.**
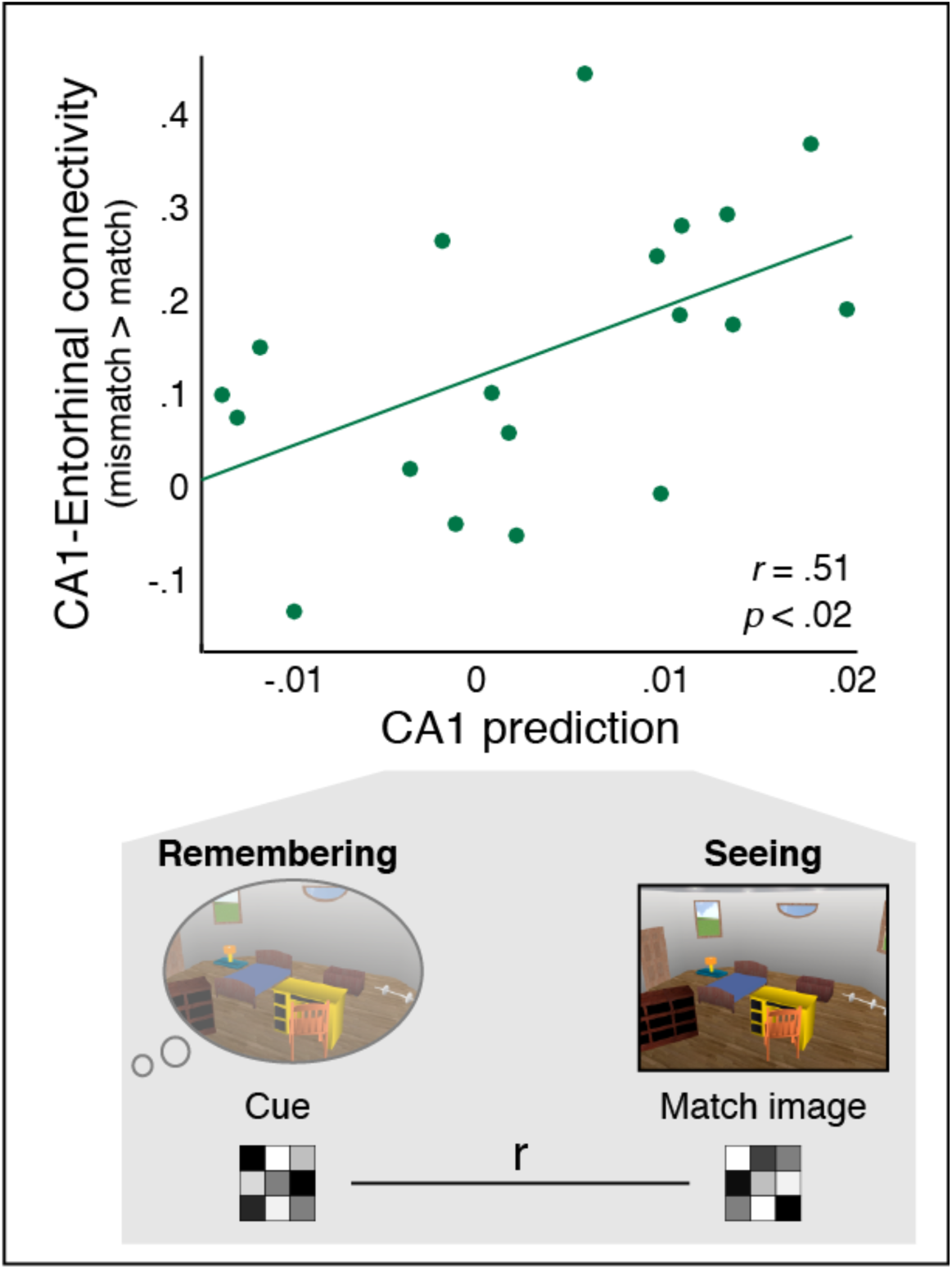
Top: CA1 prediction strength correlated with increase in functional connectivity between CA1 and Entorhinal cortex. As a connectivity measure, we took the mismatch - match contrast score for each participant (see main text). Bottom: we quantified prediction strength by computing multivariate representational similarity between the cue part of a trial, and the match image of the same room (see main text for controlling for average “room” prediction by subtracting the similarity to match images of other rooms).

### CA1 multivoxel activity patterns reflect mnemonic prediction errors

The previous analysis demonstrated that, across participants, those with stronger memory reinstatement in CA1, and presumably stronger prediction errors during viewing changes in the rooms, also had higher CA1-entorhinal connectivity in response to such violations. While the previous result addresses participants’ mnemonic *predictions*, it does not directly examine participants’ *prediction errors*. Here, we estimated a mnemonic prediction error ‘signal’ in region CA1 by measuring the difference between participants’ multivoxel activity patterns during the cue (i.e. the mnemonic prediction) and during the violations. To assess the level of mnemonic prediction errors in CA1, we computed the correlation between the multivoxel activity patterns of the prediction during the memory cue and the violation when viewing the room in the same trial. First, correlation values were submitted to a repeated-measures ANOVA, with Changes (0-4) and Task (Furniture/Layout) as within-participant factors. Since no interaction was obtained, we collapsed across tasks for further analyses (*F*(4,72) = .36, *n.s.*). We found that pattern similarity in CA1 decreased as the number of changes increased (see Figure 4). Interestingly, a match > mismatch contrast seemed to characterize the decrease slightly better than the linear contrast (match > mismatch: *t*(18) = 2.21, *p* = .04, Cohen’s *d* = .5; linear: *t*(18) = 1.54, *p* = .14, Cohen’s *d* = .35; see Figure 4). CA1 activity patterns thus are sensitive to the mismatch between a retrieved memory and perceptual input that is an altered version of that memory.

**Figure 4.**
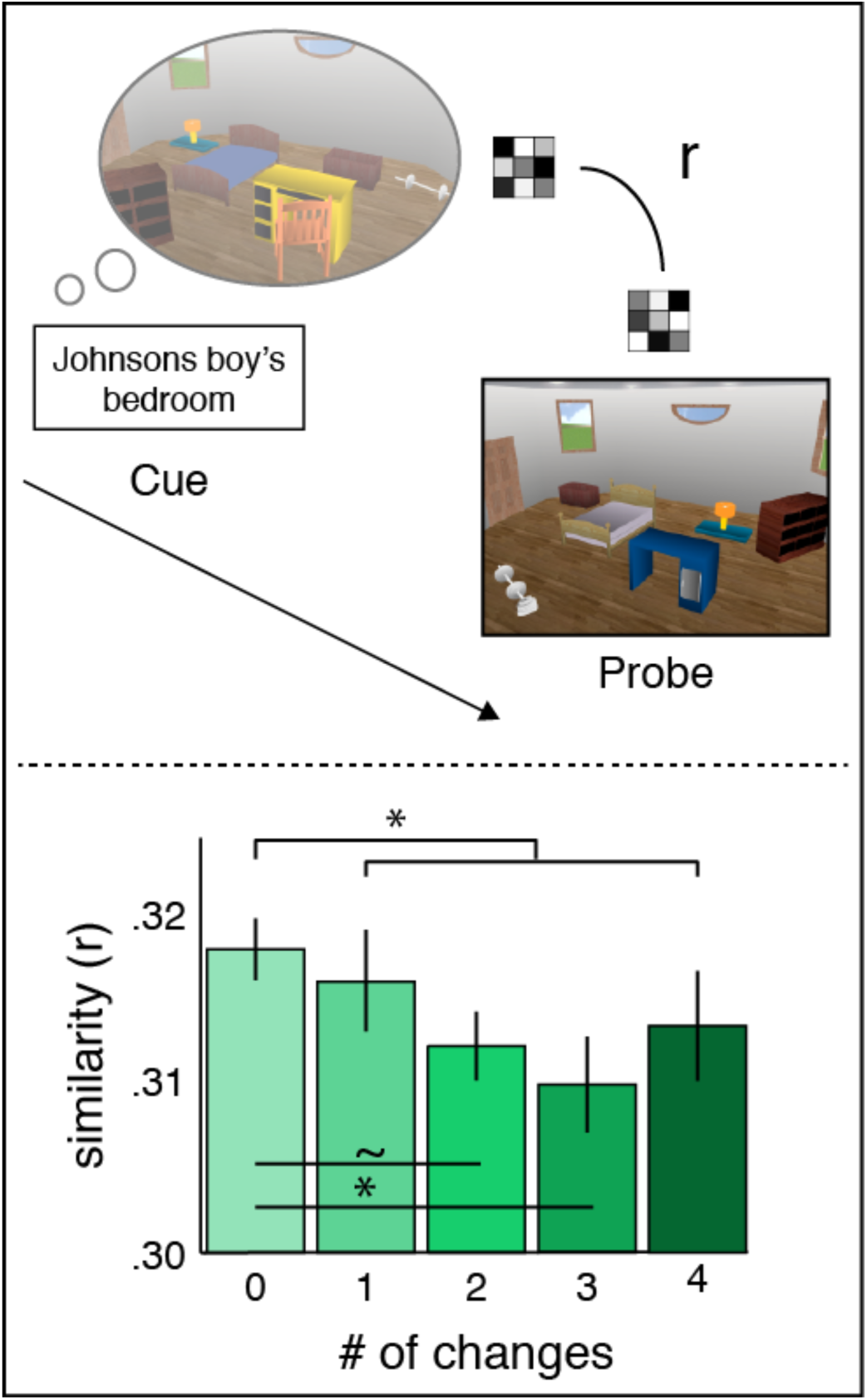
Mnemonic prediction errors in CA1. Top: Mnemonic prediction error was assessed by computing the pattern similarity between the cue and the probe parts of the trial. Bottom: CA1 similarity between the cue and the image decreased when changes were introduced in the images. * p < .05, ∼ p < .1

## Discussion

Behavioral and physiological work have implicated hippocampal processing in both laying down new memories and retrieving past memories (Eichenbaum, Yonelinas, & Ranganath, 2007; Hasselmo, Bodelón, & Wyble, 2002; Marr, 1971; O’Reilly & McClelland, 1994; Scoville & Milner, 1957; Squire & Alvarez, 1995). The computational principles that underlie these processes are in conflict as encoding will benefit most from synaptic plasticity, while, during retrieval, plasticity may alter the memory trace and lead to inaccurate memory representations (Hasselmo et al., 1996; O’Reilly & McClelland, 1994; Treves & Rolls, 1994; Yassa & Stark, 2011). To address this apparent conundrum, it has been proposed that encoding and retrieval may be mediated by distinct hippocampal ‘states’ (Colgin, 2016; Hasselmo et al., 2002; Hasselmo & Stern, 2014; Kay & Frank, 2018; Meeter et al., 2004). Specifically, recent work has linked functional coupling between CA1 and the entorhinal or perirhinal cortices with encoding and CA1-CA3 coupling with retrieval operations (Colgin et al., 2009; Duncan et al., 2014; Fernández-Ruiz et al., 2017; Hasselmo & Stern, 2014; Kemere et al., 2013; Montgomery & Buzsaki, 2007; Newman et al., 2013; Schomburg et al., 2014; Tort et al., 2009; Zheng et al., 2015).

Here we leveraged these findings to ask whether interactions between internal memory states and conflicting environmental evidence can dynamically modulate or bias hippocampal processing ‘states’ in predictable ways. To the extent that violations of expectations drive new learning or encoding, they should adaptively bias CA1 processing of inputs from medial temporal lobe (MTL) cortical regions. At the same time, these mnemonic prediction errors might down-weight projections from the now incorrect memory-based predictions from CA3 to CA1. To test that hypothesis, participants were cued to retrieve previously well-learned images of rooms. Memory retrieval was then followed by the visual presentation of images that either matched or mismatched the learned information (Methods and Results). Consistent with our hypothesis, we found that CA1 connectivity with entorhinal cortex increased as mnemonic prediction errors increased. This was accompanied by a decrease in CA1-CA3 connectivity for those same trials. Thus, mnemonic prediction errors do not simply lead to an overall general increase (or decrease) in functional connectivity of the CA1 region, but rather they selectively and differentially modulate processing along distinct hippocampal pathways.

To support the notion that connectivity changes were related to participants’ internal memory predictions, we quantified prediction strength by examining the multi-voxel similarity in CA1 between a retrieved memory of a room and viewing of the room. We found that participants with better cued memory reinstatement showed a greater increase in CA1-entorhinal connectivity in response to subsequent violations of the remembered rooms. These results suggest that an interplay between internal memory predictions and environmental evidence modulate further hippocampal processing ‘states’, potentially driving hippocampal processing towards an encoding ‘state’ and away from a ‘retrieval’ state (Colgin, 2016; Hasselmo et al., 1995, 1996; Meeter et al., 2004; O’Reilly & McClelland, 1994). Such state shifts may prove to be an adaptive mechanism for memory updating: by reducing processing of erroneous retrieved predictions while up-regulating encoding of the novel sensory evidence.

How the hippocampus shifts between memory states is largely unknown. It is possible that both acetylcholine (ACh) and dopamine (DA) play a role in biasing hippocampal states (Duncan, Sadanand, et al., 2012; Giocomo & Hasselmo, 2007; Hasselmo, 2006; Lisman & Grace, 2005; Meeter et al., 2004). Some models propose that novelty detection in the hippocampus upregulates ACh input, which in turn increases excitation in the CA1-entorhinal pathway, while dampening CA1-CA3 communication (Meeter et al., 2004; Newman et al., 2013). It has also been proposed that ACh input may further entrain theta and gamma frequencies associated with encoding versus retrieval states (Colgin et al., 2009; Hasselmo, 2006; Meeter et al., 2004; Newman et al., 2013; Vandecasteele et al., 2014). Another influential theory suggests that increased CA1 activity in response to prediction errors leads to an increase in activation in the ventral tegmental area (VTA), a primary source of DA, which in turn projects back to CA1 and entorhinal cortex (Lisman & Grace, 2005). Supporting evidence comes from fMRI studies showing concomitant hippocampal and VTA activation in response to novel and unexpected events (Bunzeck & Duezel, 2006; Wittmann, Bunzeck, Dolan, & Düzel, 2007) and VTA-CA1 interactions were recently shown to mediate associative memory encoding (Tompary, Duncan, & Davachi, 2015; see also Shohamy & Adcock, 2010, for review). In rodents, injection of DA agonist to the CA1-entorhinal pathway increased the CA1 post-synaptic potential, suggesting that DA can increase CA1-entorhinal synaptic transmission (Vago, Bevan, & Kesner, 2007; cf. Otmakhova & Lisman, 1999). Thus, it is possible that CA1 activation leads to engagement of the postulated back-projection from VTA to CA1 and entorhinal cortex (Lisman & Grace, 2005) and serves to functionally couple these regions and enhance CA1-entorhinal connectivity. Consistent with that notion, we found that connectivity in CA1-entorhinal cortex was correlated with the strength of the memory predictions measured in area CA1. Namely, those participants who showed greater similarity between a viewed room and the rooms’ retrieval cue, our measure of a mnemonic prediction, also exhibited larger increases in CA1-entorhinal connectivity in response to presented rooms that contained changes, or violations, of the learned room. More work is needed, however, to better understand how neurotransmitters such as DA, ACh, and potentially norepinephrine (Clewett, Huang, Velasco, Lee, & Mather, 2018; Giocomo & Hasselmo, 2007; Kafkas & Montaldi, 2018) contribute to a shift in hippocampal connectivity with changing mnemonic demands.

While the accounts discussed above place the CA1 region as the source of violation-detection and connectivity changes (Chen et al., 2015; Duncan, Ketz, et al., 2012; Hasselmo et al., 2002, 1996; Kumaran & Maguire, 2007b, 2009; Lisman & Grace, 2005; Meeter et al., 2004), it is possible that prediction errors are also detected in earlier brain regions. Experimental and computational work in the predictive coding framework converge on the notion that high-level areas project top-down predictions to earlier visual cortices, where these predictions are then compared to incoming sensory information (Friston, 2005; Friston, 2018; Rao & Ballard, 1999). Consistent with this, after learning that a stimulus predicts another visual stimulus, greater activity was reported in visual cortex of both humans and monkeys in response to stimuli that violated such memory-based predictions, compared to stimuli that confirmed prior expectations (e.g. Kok, Jehee, & de Lange, 2012; Meyer & Olson, 2011: for recent reviews, see e.g., de Lange, Heilbron, & Kok, 2018; Ouden, Kok, & Floris, 2012). Moreover, it is now widely reported that memory reinstatement in cortical regions is correlated with hippocampal activity (Bosch, Jehee, Fernandez, & Doeller, 2014; Danker, Tompary, & Davachi, 2017; Hindy, Ng, & Turk-Browne, 2016; Kok & Turk-Browne, 2018; Long, Lee, & Kuhl, 2016; Ritchey, Wing, Labar, & Cabeza, 2012; Staresina, Henson, Kriegeskorte, & Alink, 2012). While fMRI studies cannot resolve the temporality of neural activity, a recent ECoG study found that memory reinstatement in visual processing regions preceded hippocampal reinstatement in humans (Lohnas et al., 2018). Together, these studies suggest that memory reinstatement, or predictions, may occur in early processing stages, and hence then influence subsequent hippocampal processing. Like memory-predictions, it is also possible that early prediction-error signals as those mentioned above may propagate forward to influence hippocampal processing (Henson & Gagnepain, 2010) and potentially mediate connectivity changes.

A critical assumption in models of CA1 function is that CA1 may be ideally suited to compare internal memory output with input from visual cortical regions representing ongoing visual experience (Hasselmo & Wyble, 1997; Hasselmo et al., 1996; Kumaran & Maguire, 2007b; Lisman & Grace, 2005). While earlier investigations have reported increased BOLD signal during mnemonic prediction errors in the hippocampus and, specifically, in area CA1 (Chen et al., 2015; Duncan, Ketz, et al., 2012; Kumaran & Maguire, 2006, 2007a), these studies did not specifically measure memory predictions in CA1, nor could they address the content of CA1 processing. Thus, whether the content of CA1 processing indeed reflects predictions as well as incoming sensory input, or whether univariate findings reflect other violation-related processes remained unknown. Here, we found that in CA1, activity patterns during cued memory reinstatement were more similar to activity patterns during viewing the same image, compared to viewing an altered version of image (Results, Figure 4). This result suggests that the content of CA1 representations are sensitive to the difference between internal memory representations and sensory evidence, thus providing essential evidence to support the role of CA1 as a violation detector (Hasselmo & Wyble, 1997; Hasselmo et al., 1996; Kumaran & Maguire, 2007b, 2009; Lisman & Grace, 2005).

In summary, we found that mnemonic prediction errors biased hippocampal area CA1 connectivity towards entorhinal cortex and away from area CA3. We propose that this bias may reflect a shift in hippocampal ‘states’ towards encoding of the novel sensory information and away from retrieval of erroneous memory-based predictions. How the hippocampus supports both encoding and retrieval is an intriguing question that has received increased attention in recent years (Colgin, 2016; Colgin & Moser, 2010; Duncan, Sadanand, et al., 2012; Hasselmo & Stern, 2014). The current results contribute to this on-going line of research by measuring hippocampal states in humans, and by suggesting that the interplay between memory reinstatement as a prediction and their subsequent violation, or mnemonic prediction errors, may be an important factor in biasing these states. Thus, in addition to understanding the distinct neural mechanisms that allow shifting between encoding and retrieval, future research should aim at understanding the psychological factors that may shift our cognitive system between these different mnemonic states (Duncan, Sadanand, et al., 2012; Hasselmo et al., 2002; Meeter et al., 2004).

## Author contributions

O.B. and L.D. and KD conceptualized the general experimental design. O.B. and L.D. conceptualized the specific analytic approach reported in this manuscript. O.B. analyzed the data. K.D. conducted the preprocessing of fMRI data and regions of interest demarcation. O.B., L.D. and K.D. wrote the paper. K.D. and L.D. conceptualized and designed the task. K.D. collected the data.

## Acknowledgements

This research was supported by the National Institute of Mental Health Grant R01MH074692 to L.D. O.B. is further supported by McCracken fellowship.

## Methods

### Participants

Twenty participants were included in the current study (Mean age: 25.4 years). Further information can be found in Duncan et al. (2012), where the results of univariate analyses of these data were previously published. One participant was removed from all analyses due to substantial entorhinal dropout (see Regions of Interest).

### Procedure

In the training phase (∼24h prior to scanning, and again before entering the scanner), participants were extensively trained to identify each of 30 named rooms (e.g., “Johnson’s boy bedroom”) to criteria (Duncan et al., 2012). While scanning, participants were employed in two change-detection tasks. In both tasks, the room’s name appeared for 1.5 s, followed by 1 s blank and a probe image (4 s). The probe image contained 0-2 changes in the individual pieces of furniture, along with 0-2 changes in the layout of the furniture, relative to the learned image, making a total of 0-4 changes per image. In the Furniture task, participants were asked to indicate whether all pieces of furniture were identical to the studied image. In the Layout task, participants were asked to indicate whether the layout of the furniture was identical to the learned image. This resulted in a 2 (Task: Furniture/Layout) by 5 (Changes: 0-4 total changes) within-participant design. Each room appeared once in every trial type (9 trial types: 0/1/2 furniture changes by 0/1/2 layout changes), across both tasks, to make a total of 270 trials. Here, we focused on total number of changes (0-4 total changes, see below). Thus, analysis was conducted on 30 trials in each of the 0 and 4 changes, 60 trials in the 1 and 3 changes conditions, and 90 trials in the 2 changes condition (across both tasks). Tasks were blocked, such that each scan included one task (10 scans, 5 per task), and the blocks alternated between the Furniture and the Layout task. One participant had 8 blocks, and another had 7. The minimal number of trials per condition was 24 and 21, correspondingly, still allowing a meaningful analysis. Hence these participants were included in the analysis.

### FMRI parameters

Scanning was performed using a 3T Siemens Allegra MRI system. A high-resolution EPI sequence was used to collect functional data (TR=2.5 s, TE=49 ms, FOV = 192 × 96 mm, 26 interleaved slices, distance factor of 20%, 1.5 × 1.5 × 2 mm voxel size). A T1-weighted high-resolution MPRAGE (1 × 1 × 1 mm voxel size) was used as an anatomical scan.

### Regions of Interest (ROIs)

Anatomical ROIs were drawn manually by K.D. on each participant’s MPRAGE anatomical image, and were then registered to functional space. The same hippocampal ROIs (CA1, CA2/CA3/DG) reported in Duncan et al. (2012) were used here. These ROIs were drawn in a similar procedure to Kirwan et al. 2007 (Duncan, Ketz, et al., 2012; Kirwan, Jones, Miller, & Stark, 2007). The entorhinal cortex was drawn using guidelines discussed by (Insausti et al., 1998; Pruessner et al., 2002). ROIs were also masked to remove voxels with substantial signal dropout, a concern mainly in the entorhinal cortex (Carr, Rissman, & Wagner, 2010). One participant with only 12 voxels in the left entorhinal and 80 voxels in the right entorhinal was excluded from all analyses. All other participants had on average 234 voxels in the left entorhinal ROI (range: 127-344), comprising 84% (range: 44%-93%) of the anatomical left entorhinal. In the right entorhinal ROI, participants averaged 255 (range: 165-337) voxels, which were 87% (59%-95%) of the anatomical right entorhinal.

### Functional connectivity: fMRI beta-series correlation

Functional connectivity between regions was computed using a beta-series correlation approach (Rissman et al., 2004), in which a timeseries of single-trial parameter estimates in two regions are correlated. To obtain the single-trial estimates we used an LSS (Least-Square-Separate) approach (Mumford, Davis, & Poldrack, 2014; Mumford, Turner, Ashby, & Poldrack, 2012; Turner, Mumford, Poldrack, & Ashby, 2012). We reasoned that this approach would maximize our ability to capture the variance explained by the image portion of each trial (our focus of interest) and distinguish this variance from preceding cue part of each trial (the name of each room). Thus, in the first level analysis, a separate GLM was computed for each trial. Each model included the image portion of a single trial as a regressor of interest. The cue portion in all trials were included in one regressor of no interest. Other images were binned based on trial type to make 9 additional regressors of no interest. In all regressors, events were modeled as boxcars lasting for the duration of the event (1.5s for cues, 4s for images) convolved with a double gamma function to approximate the hemodynamic response. A temporal derivative regressor was also added for each regressor. GLMs were implemented using FSL FEAT. This procedure yielded 270 parameter estimates, one for each trial. A t-stat was computed for each parameter estimate, and these were averaged, per each trial, across all voxels in each ROI (CA1, CA2/CA3/DG, entorhinal cortex, and perirhinal cortex, separately for right and left hemispheres). T-stats were then binned based on experimental conditions: number of changes (0-4) and task (Furniture/Layout) to make 10 t-series for each ROI. We then computed functional connectivity between area CA1 and the other brain regions of interest: CA2/3/DG and entorhinal cortex in each of the 10 conditions separately for each hemisphere. The Pearson’s r values per each participant, condition and pair of ROIs were Fisher transformed and entered to the group-level analysis.

### CA1 mnemonic prediction strength analysis

In order to measure the strength of participants’ mnemonic predictions, we used a representational similarity analysis (RSA; Kriegeskorte, Goebel, & Bandettini, 2006; Kriegeskorte, Mur, & Bandettini, 2008). To obtain the multivoxel activity pattern for each cue, we used the same LSS procedure as for the images (see Functional connectivity: beta-series correlation). Each cue was allocated a separate GLM, which included one regressor of interest for the cue, and a few regressors of no interest: one regressor for all other cues, and 9 additional regressors modelling the images – one for every trial type. As with the image models, a time-derivative regressor was added for each regressor. Parameter estimates were then converted to t-statistics, which were taken to the RSA.

To compute the strength of participants’ mnemonic predictions, we correlated the multivoxel activity pattern in CA1 observed in response to each room cue with the multivoxel activity pattern measured when participants viewed the intact room image (i.e., the 0-changes image). For example, the CA1 activity pattern in response to the verbal cue “Johnsons boy’s bedroom” was correlated with the CA1 activity in response to the intact image of Johnsons boy’s bedroom. To compute the similarity to the specific match image, while controlling for condition-level effects and general similarity to all 0-changes images, we computed, for each cue, the correlation between the activity pattern during the cue and the activity pattern of other 0-changes images, and averaged across these correlation values. Then, we subtracted this average correlation with other 0-changes images from the correlation with the intact image corresponding to the cue (e.g., the intact image of Johnsons boy’s bedroom). This yielded, for each cue, a measure of how good the prediction of the specific corresponding room was, beyond overall similarity to a 0-changes image. This procedure further controlled for differences in average similarity values between participants, which is critical for a meaningful interpretation of across participant correlations of prediction strength with connectivity. Cues in some trials were excluded from this analysis: first, we excluded cues in the 0-changes condition. These cues were presented in the same trial as the corresponding intact image while all other 0-changes images were presented in other trials, thus we avoided comparing within-trial similarity to across-trial similarity. Second, we excluded cues and intact images that were presented in the same scan to avoid inflating similarity values within the same scan (Mumford et al., 2014). Third, we only took cues in which the cue and the intact image were presented in the same task, to avoid introducing task differences between the cue and the image. For each participant, the correlation values between the cues that entered the analysis and their corresponding 0-changes images (other 0-changes images subtracted, as detailed above) were averaged and Fisher-transformed to obtain a prediction index per participant. These values were then used to correlate the prediction strength with CA1-entorhinal connectivity. As a connectivity measure summarizing the change in connectivity in response to mnemonic prediction errors per participant, we used the match < mismatch contrast score, computed by multiplying, per participant, the connectivity in the 0-changes condition by −1, and each of the number of changes (1-4) by .25, and summing these values (see also below). This contrast revealed to well characterize CA1-entorhinal connectivity (see Results).

### CA1 multivariate mnemonic prediction error analysis

To further support our hypothesis that mnemonic prediction errors modulate hippocampal connectivity, we aimed to compute a measure of mnemonic prediction error in our study. To this end, we correlated the CA1 activity pattern during the presentation of each cue when participants were instructed to retrieve a memory of the cued room (i.e., the mnemonic prediction) with the CA1 activity pattern measured when viewing the probe image on each trial (the sensory evidence). We reasoned that the difference between the representation of the mnemonic prediction and that of the sensory evidence can be interpreted as mnemonic prediction error. We averaged this value across all the trials within each number of changes (0-4), and separately in each task, and Fisher-transformed these correlation values for statistical analysis. If indeed participants retrieved the intact image on each trial, we predicted a decrease in similarity, or increased prediction error, as number of changes increased, reflecting larger divergence between the retrieved memory and the sensory evidence.

### Statistical tests for the functional connectivity analysis

In the group-level analysis of the functional connectivity data (beta-series correlation), Fisher-transformed r values in each pair of ROIs were entered to a 5 (Changes: 0-4) by 2 (Task: Furniture/Layout) repeated-measures ANOVA. We saw no interaction between Task and Changes in CA1 connectivity with either CA3 or entorhinal cortex. Thus, for CA1-CA3 and CA1-entorhinal, for each participant in each number of changes, we collapsed across tasks to obtain an average beta-series correlation value. To directly test our hypothesis that mnemonic prediction errors modulate CA1 connectivity with CA3 vs. entorhinal cortex, we conducted a 5 (Changes: 0-4) by 2 (ROI: CA3 vs. entorhinal) repeated-measures ANOVA. Where a Changes by ROI interaction was observed, we tested how Changes (0-4) influenced connectivity separately in each pair of ROIs (CA1-CA3, CA1-entorhinal), using a one-way repeated-measures ANOVA.

Although we had no specific hypothesis regarding the shape of the increase or decrease in connectivity, we sought to further characterize connectivity changes. We asked whether connectivity changed linearly with number of changes, or, alternatively, whether changes may reflect a binary match-mismatch signal, whereby any level of change is different from no-changes at all, with no or little difference between level of changes. To that end, we defined a linear contrast by allocating for each number of changes (0,1,2,3,4) linear-trend values (−2,−1,0,1,2) correspondingly. The match < mismatch contrast was defined as by coding the 0-changes condition as −1, whereas the 1-4 changes conditions were coded 0.25 each. We directly compared the linear trend contrast to the match < mismatch contrast by using a mixed-effects model approach as implemented by lmer function in R (Bates, Mächler, Bolker, & Walker, 2014). We included both contrasts as explanatory variables in the same model (the beta series correlation value per participant per number of changes was the explained variable) and then compared this full model to either a model including only the linear trend contrast, or only the match < mismatch contrast (match < mismatch was treated as a factor, an intercept per participant was included in all models). This analysis thus examines whether one contrast significantly explains variance above and beyond the other contrast.

### Statistical tests for the prediction strength and mnemonic prediction error analyses

The significance of the correlation of prediction strength with functional connectivity was tested using a one-tailed t-test for Pearson’s correlation. One-tailed was used since there was a clear prediction that stronger predictions would correlate with connectivity changes.

For the mnemonic prediction error analysis, we first entered the Fisher-transformed similarity values to a 5 (Changes: 0-4) by 2 (Task: Furniture/Layout) repeated-measures ANOVA. To preview, since there was no interaction between Changes and Task in CA1, we collapsed across Task in all further analyses. Like in the functional connectivity analysis, we then estimated this decrease using a linear trend analysis, as well as a match < mismatch analysis, using the same contrasts as described above.

